# Stability in the genetic structure of a *Zymoseptoria tritici* population from epidemic to interepidemic stages at a small spatial scale

**DOI:** 10.1101/222380

**Authors:** D. Morais, C. Duplaix, I. Sache, V. Laval, F. Suffert, A-S. Walker

## Abstract

Subpopulations of the wheat pathogen *Zymoseptoria tritici* (26 sample groups composed of 794 strains) were collected in two nearby wheat fields in the Paris basin, during both epidemic and inter-epidemic periods of three successive years (2009-2013). In addition to the type of inoculum (ascospores *vs*. pycnidiospores), the alternative presence of wheat debris allowed taking into account its putative origin (local *vs*. distant). We used a molecular epidemiology approach, based on population genetic indices derived from SSR marker analysis, to describe putative changes in the structure and genotypic diversity of these subpopulations over three years, at a spatiotemporal scale consistent with epidemiological observations. Genetic structure was stable over time (within and between years) and between fields. All subpopulations displayed very high levels of gene and genotypic diversity. The low levels of linkage disequilibrium and the very low clonal fraction at all stages were consistent with the regular occurrence of sexual reproduction in the two fields. A significant increase of the MAT1-1/MAT1-2 ratio was observed over the course of the epidemics, suggesting a competitive advantage of MAT1-1 strains consistently with their greater pathogenicity reported in the literature. Finally, we found that the period, the type of inoculum and its putative origin had little effect on the short term evolution of the local population of *Z. tritici*. Fungal population size and diversity are apparently large enough to prevent genetic drift at this fine spatiotemporal scale, and more likely short distance migration contributes strongly to the stabilization of genetic diversity among and within plots.

## Introduction

Septoria tritici blotch can cause yield losses of 10-40% (HGCA, 2012) on wheat (*Triticum aestivum*) crops in several wheat-growing areas worldwide. This disease is caused by the hemibiotrophic, heterothallic, ascomycete fungus *Zymoseptoria tritici*. The means of disease management include fungicide sprays, the use of tolerant cultivars, and cultural practices such as crop rotation and stubble management that decrease the amount of primary inoculum (Eyal, 1999). Ascospores ejected from contaminated debris from the previous wheat crop (Sanderson, 1972) and dispersed over long distances by the wind, are the principal source of primary inoculum for *Z. tritici* infections. By contrast, pycnidiospores, which are splash-dispersed over shorter distances by the rain, are the main source of secondary inoculum during the growing period of the wheat crop (Shaw & Royle, 1989; Suffert *et al*., 2011). Furthermore, ascospores and pycnidiospores may both be involved in epidemic development, albeit at different stages (Suffert & Sache, 2011). In a wheat monoculture system in which contaminated debris is not completely buried, pycnidiospores can, theoretically, contribute to initiation of the epidemic (Suffert *et al*., 2011). In the spring, secondary infections may also be induced by ascospores released from old, still infectious debris in distant plots or from infected wheat plants located in the same plot (Duvivier, 2015). The level of sexual recombination in *Z. tritici* populations and the relative contributions of different types of inoculum (pycnidiospores *vs*. ascospores; Zhan *et al*., 1998) to the epidemic development are all the more difficult to determine experimentally since they depend on the year climatic conditions (Duvivier, 2015). Simulations with a mathematical model including the effect of spore type on epidemic development suggested that sexual reproduction can occur and affect the genetic composition of the population during the growing season (Eriksen *et al*., 2001). Suffert & Sache (2011) and Morais *et al*. (2016a) showed that an epidemic developed more rapidly in a plot containing ascospore-releasing debris than in a plot without debris. Morais *et al*. (2016b) also showed how the differential adaptation of a resident population (i.e. originating from wheat debris of a monoculture plot) to local *vs*. distant host cultivar, can be used to infer the origin of primary inoculum. Altogether, these various observations highlight multiple and complex contributions of inoculum sources on the onset of Septoria tritici blotch epidemics and the subsequent development of *Z. tritici* populations. Approaches complementary to classical epidemiology would be needed to disentangle the contributions of the various phenomena at work.

The spatial patterns and genotypic diversity of pathogen populations could be useful for inferring the type of inoculum playing the most important role during an epidemic period (Milgroom & Peever, 2003). Observations and modeling of the spatiotemporal dynamics of the propagules thought to act as primary inoculum can lead to the identification of sources of primary inoculum (Rieux *et al*., 2014). Population genetics also provides information about the evolutionary processes involved in the establishment and maintenance of genetic variation within and between populations, by focusing on changes in multilocus genotypes or allele frequencies (McDonald & Linde, 2002; Abang *et al*., 2006). Population genetics studies could, therefore, be used to determine when the primary inoculum is produced and its contribution to the onset of the epidemic (Zwankhuizen *et al*., 1998; Shah *et al*., 2001; Milgroom & Peever, 2003). In this approach, the primary inoculum may be identified as new genotypes produced through sexual reproduction and/or airborne propagules of remote origin, which theoretically may modify genotypic diversity and clonal fractions of the population.

Empirical investigations of the diversity and structure of populations of *Z. tritici* on a single date during annual epidemics have mostly been carried out at a large geographic scale (country to continent; Zhan *et al*., 2002; 2003; Razavi & Hughes, 2004; Abrinbana *et al*., 2010; El Chartouni *et al*., 2011; Boukef *et al*., 2012), with only a few rare studies focusing on local (field; Zhan *et al*., 2001; El Chartouni *et al*., 2012; Siah *et al*., 2018) or very local (plant or leaf; Linde *et al*., 2002) scales and involving assessments at multiple time points in the epidemic stage (Chen *et al*., 1994). Despite potential differences in the type and amount of inoculum at different time points within an epidemic, the period of *Z. tritici* population sampling has rarely been considered as a factor potentially affecting genetic structure. Recent epidemiological studies suggest that short-term evolution processes – including selection and counter-selection – can be driven by host cultivar adaptation and result in a trade-off between intra- and interannual scales which can be phenotypically detectable (Suffert *et al*., 2015; 2018). *Z. tritici* populations have been reported to display very high levels of genetic diversity and no structure at the continental and regional scales (Schnieder *et al*., 2001; Linde *et al*., 2002), except in the studies of El Chartouni *et al*. (2011; 2012), Abrinbana *et al*. (2010) and Siah *et al*. (2018). Relatively little is ultimately known about the evolution of *Z. tritici* populations over a fine spatiotemporal scale, within and among epidemic seasons. Such knowledge could be of particular interest to investigate the short-term dynamics of *Z. tritici*, particularly the adaptation of the pathogen population (virulence and aggressiveness) to the host resistance. We could hypothesize that changes in diversity occurs locally at the year scale and is driven by changes in the relative importance of different contamination processes during epidemics. At the early stages of an epidemic the initial (founder) population consisted of ascospore-derived strains having a more or less distant origin (depending on the local presence of wheat debris) and so having mainly a non-local evolutionary history. Dispersal of ascospores leads to allele pool homogenization and dispersion of new genotypes at small spatial scales including field (Linde *et al*., 2002). Conversely, at the late stages of epidemic, the part of the population responsible for secondary infection consisted of strains having a local origin; this population is mainly derived from asexual reproduction and was potentially the result of a seasonal short-term selection driven by abiotic (fungicides sprays; temperature and host stage; Suffert *et al*., 2015) and biotic factors (host resistance; Morais *et al*. 2016b).

In this study we developed a molecular epidemiology approach to compare the structure of populations in fields with and without inoculum production *in situ* from crop debris among three successive epidemics and to identify possible structuring factors. We deliberately focused on a fine spatiotemporal scale since the monoculture field scale offers the possibility to avoid confounding effects (*e.g*. from the cultivars, fungicide treatments or cultural practices). We assessed and analyzed changes in neutral structure and diversity of several subpopulations collected in two nearby wheat fields during both epidemic and interepidemic periods of three successive years (2009-2013). The presence of wheat debris in one of the fields, yearly, enabled to contrast them for the origin of the inoculum (resident, constituted of local ascospores and pycnidiospores *vs*. distant, constituted of wind-dispersed ascospores from other fields). As it was already shown in the same study area that the local presence of inoculum had a strong quantitative effect, although transient, on the early epidemic dynamics (Suffert & Sache, 2011; Morais et *al*., 2016a), we cannot exclude consequences on the genotypic composition of the pathogen population.

## Materials and methods

### Experimental design and sampling

A four-year field experiment was carried out from 2009 to 2013 at the INRA Grignon experimental station (France, 48°51’N, 1°58’E), on the basis of the field design described by Morais *et al*. (2016b). Winter wheat (cv. Soissons) was grown as a monoculture on a 20 × 100 m plot (D+) during the 2009-2010, 2010-2011, 2011-2012 and 2012-2013 cropping seasons, with no fungicide applications since 2007. The debris from the previous wheat crop was left on the soil surface and so could act as a local source of primary inoculum. Another 40 × 100 m plot (D_0_; different location each year but 0-300 m distant from the D+ plot) on which wheat had not been grown in the preceding year, was planted with wheat in each of the four seasons. The primary inoculum infecting this plot was, therefore, assumed to have a distant origin. The D+ and D0 plots were contiguous in the 2009-2010 and 2011-2012 seasons, but were located 300 m apart during the 2010-2011 and 2012-2013 season.

Sample groups originating from pycnidia (hereafter called “pycnidiospore-derived subpopulations”) were simultaneously collected from wheat leaves infected with *Z. tritici* in the D+ and D_0_ plots, at one-to two-month intervals, from October 2009 to December 2012. Fungal isolates were obtained by placing the infected leaves in a plastic box with wet paper and incubating them overnight at 18°C, to promote cirrhus exudation. We removed one cirrhus per pycnidium and per leaf lesion for culture in a single Petri dish filled with PDA medium (potato dextrose agar, 39 g.L^−1^) for five days. We obtained 681 pure isolates (Fig. 1). Samples collected on dates that were close together were grouped together by epidemic stage, to constitute 26 representative subpopulations of 16 to 84 individuals (P1 to P29). Consequently, four groups were studied for the 2010-2011 and 2011-2012 cropping seasons (Fig. 1): “beginning of the epidemic period” (from October to January, corresponding to the fall and the first few weeks of winter, when the wheat was starting to tiller; subpopulations P02, P05, P09, P13, P17, P23, and P29), “intermediate epidemic period 1” (from February to late March, corresponding to the end of winter and the beginning of spring; subpopulations P10, P14, P18, and P24), “intermediate epidemic period 2” (from April to May, corresponding to the spring; subpopulations P03, P06, P11, P15, P19, and P25) and “end of the epidemic period” (from late-May to mid-July, corresponding to the end of spring and the first few weeks of summer, when the wheat reached the mature adult stage; subpopulations P04, P07, P12, P16, P21, and P27). For the 2009-2010 season, only three groups were studied due to the low disease severity in the “intermediate period”, for which sampling was carried out on only one date in March 2010, the “beginning period” and the “end period” being similar, as previously described.

**Figure 1:**
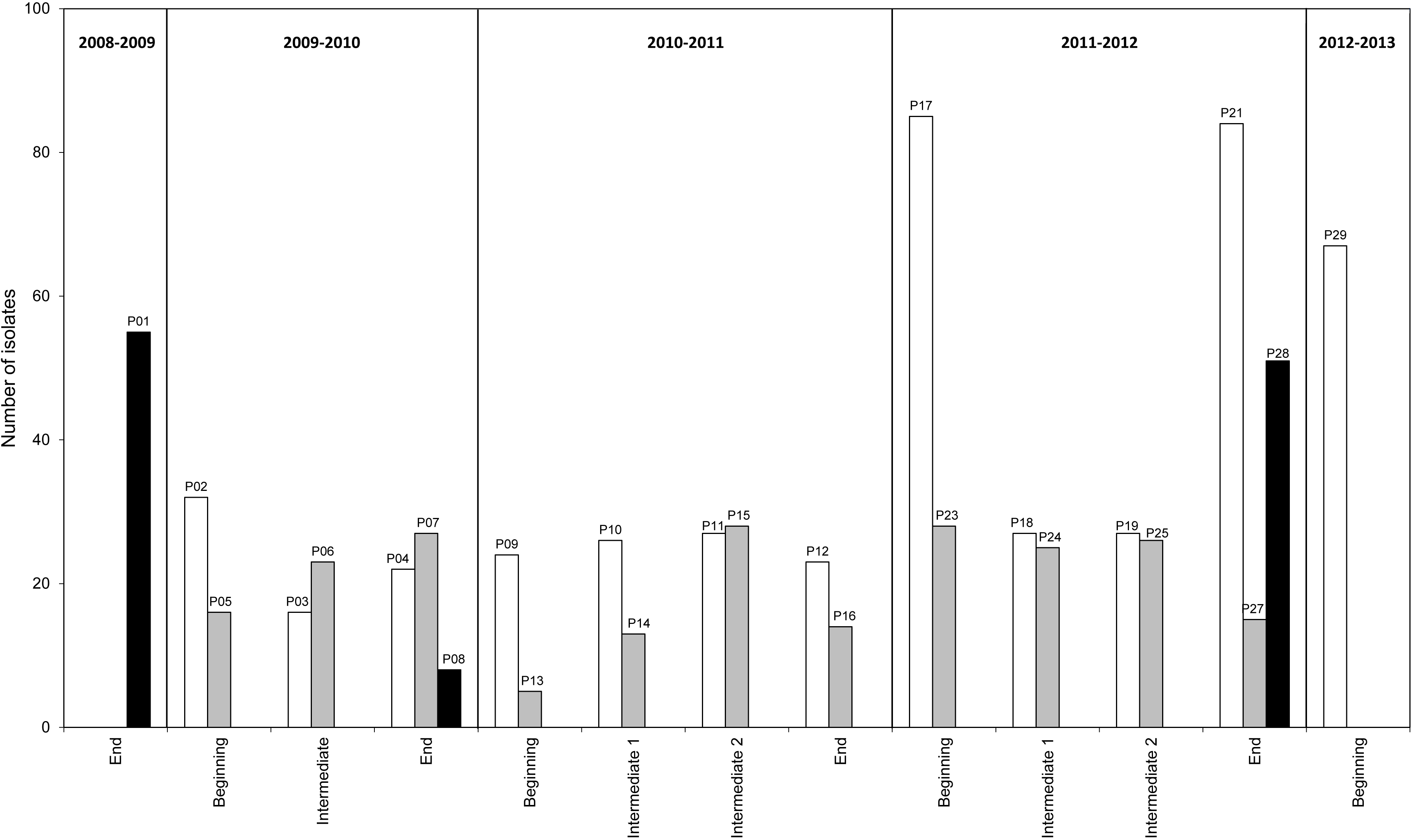
Subpopulations of *Z. tritici* (794 isolates) collected from two wheat plots (D+, with debris; D_0_, without debris) over the course of three successive annual epidemics (2009-2010, 2010-2011 and 2011-2012) in Thiverval-Grignon, France. The 794 isolates were grouped according to the epidemic period (beginning, i.e. from late-November to late-January; intermediate 1, i.e. February and March; intermediate 2, i.e. from April and May; end, i.e. from early-June to mid-July). White columns: pycnidiospore-derived subpopulations from the plot with debris (D+); gray columns: pycnidiospore-derived subpopulations from the plot without debris (D_0_); black columns: ascospore-derived subpopulations from the plot with debris (D+).

Sample groups originating from pseudothecia (hereafter called “ascospore-derived subpopulations”) were collected after the interepidemic period, from wheat debris bearing pseudothecia in the D+ plot. Wheat debris was kept in damp plastic boxes at 18°C overnight. The next day, the debris was cut into small pieces, each lot weighing 10 g, and spread out in fresh damp boxes (24 × 36 cm). Eight Petri dishes filled with PDA medium were placed upside down 1 cm above the debris in each box. The damp boxes were then incubated in the dark for at least 6 h at 18°C before being covered and incubated in the same conditions until the formation of colonies, which were then isolated. We obtained 115 isolates corresponding to three subpopulations (P01, P08, and P28).

The 794 pure strains used in this study were stored at −80°C in a 1:1 glycerol/water solution.

### Molecular procedures

For each isolate, DNA was extracted from 50 mg of fresh fungal material scraped from a five-day culture grown on PDA medium in the dark at 18°C. DNA was extracted in an automated system (Biorobot 3000, Qiagen), with the appropriate DNeasy kit (Qiagen). All 794 isolates were genotyped for 11 single sequence repeat (SSR) markers (St2, St3A, St3C, St4, St5, St6, St7, St9, St10, St11 and St13; Gautier *et al*., 2014) in two multiplex PCRs, ensuring that amplicons with similar magnitudes of sizes bore distinct fluorochromes. Automatic allele recognition and annotation, based on binning analysis, were performed, to ensure the homogeneous naming of the alleles at each locus (Beckmann Coulter CEQ 8000 software; Gautier *et al*., 2014). In total, 585 isolates were successfully genotyped with the 11 SSRs. The remaining isolates (27% of the total dataset) provided no amplicons for only one or two markers, even spread between the SSRs; they were treated as missing values in subsequent analysis. Because the proportion of identical genotypes (clones) in the subpopulations was very low, genetic analyses were performed without clone correction.

Mating type was determined by PCR for 739 isolates (Waalwijk *et al*., 2002). Eight per cent of the isolates were not genotyped for mating type due to technical failures or the contamination of PCR assays.

### Analyses of population subdivision without *a priori* assumptions

Population subdivision without *a priori* assumptions was investigated by the Bayesian clustering method implemented in the *Structure* program (Pritchard *et al*., 2000). This model-based algorithm assumes linkage equilibrium within the inferred genetic clusters, and is, therefore, appropriate, in principle, for species in which recombination is a regular occurrence. We run *Structure* with the admixture model and correlated allele frequencies. The burn-in length was set at 500,000 Markov Chain Monte-Carlo iterations. The burn-in period was followed by a run phase of 1,000,000 iterations (El Chartouni *et al*., 2011), with the number of clusters *K* ranging from 1 to 10, and 10 independent replicates for each value of *K*. The amount of additional information explained by increasing *K* was determined by calculating the Δ*K* statistic (Evanno *et al*., 2005) with *Structure Harvester* (Earl & von Holdt, 2012; Belkhir *et al*, 1996-2004).

Three *Structure* analyses were carried out independently, to investigate three factors thought likely to affect the structure of the dataset: 1) collection time, during one of three or four epidemic periods during the three cropping seasons, 2) isolate origin (plot with or without debris), and 3) spore type giving rise to the isolate (ascospore or pycnidiospore). A first set of analyses was carried out to assess, for each cropping season separately (2009-2010, 2010-2011, 2011-2012), the effect of epidemic period on the structure of the pycnidiospore-derived subpopulations collected from the plot with debris (P02-P03-P04, P09-P10-P11-P12, and P17-P18-P19-P21, respectively). An analysis grouping together the three cropping season datasets plus the data for P29 (2012-2013), was also carried out to assess the effect of year. A second set of analyses was carried out to assess, for each cropping season separately (and then overall), the effect of isolate origin (plot with or without debris) on the structure of pycnidiospore-derived subpopulations (P02-P03-P04-P05-P06-P07, P09-P10-P11-P12-P13-P14-P15-P16, and P17-P18-P19-P21-P23-P24-P25-P27, respectively). Finally, we also analyzed the structuring effect of spore type giving rise to the isolate on the complete dataset (combining P01-P08-P28).

### Analyses of population subdivision with *a priori* assumptions

Pairwise Weir & Cockerham’s *F_ST_* index values (Weir & Cockerham, 1984) were calculated between pairs of populations defined in Fig. 1, with *Genepop* V4.1 (Raymond & Rousset, 1995). The unbiased fixation index *FST* is widely used to assess the degree of genetic differentiation between populations. It can be viewed as the proportion of genetic diversity due to allele frequency differences between populations or as the correlation between alleles within populations relative to the entire population. The significance of *F_ST_* values was calculated after 1000 randomizations. Bonferroni correction was applied to determine the significance of *P* values.

Hierarchical analyses of molecular variance (AMOVA) were performed with *Arlequin V3.5* (Excoffier & Lischer, 2010) to investigate the relative contributions of time (year and epidemic period), isolate origin and spore type giving rise to the isolate, to the partitioning of genetic variance. Each factor was tested individually with *a priori* subpopulations defined in Fig. 1 and one-or two-factor AMOVAs.

### Genetic variability and occurrence of sexual reproduction

*Genetix* (Belkhir *et al*., 1996-2004) was used to estimate within-population genetic variability over time from gene diversity, calculated as multilocus observed heterozygosity (*H_E_*), and allele richness (Ar, mean number of alleles per locus), in all populations. Paired-sample t-tests were used to assess the significance of differences between the D+ and D_0_ plots over time (with an alpha risk of 5%).

We determine the number of unique multilocus genotypes (G) and the derived clonal fraction (1-G/N) with *Multilocus* V1.3b (Agapow & Burt, 2001). This indexes use genotypic diversity to detect indirect evidence of sexual reproduction in *Z. tritici* populations over three cropping seasons. This software was also used to estimate multilocus linkage disequilibrium over time, by calculating the 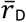 index, which is corrected for the number of loci considered, and takes values from 0 (all individuals genetically different, and multilocus linkage is broken) to 1 (all individuals are clones). The significance of the 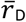 values obtained was established by comparing the observed values with the distributions obtained for 1000 randomizations (Agapow & Burt, 2001). Finally, we assessed the likelihood of recombination by sexual reproduction by calculating the ratio of mating-type alleles (MAT1-1/MAT1-2). χ^2^ tests were performed to determine whether the frequencies of the two mating types within different populations departed from the null hypothesis of a 1:1 ratio. We performed paired-sample *t*-tests to identify significant differences over time between the D+ and D_0_ plots (alpha risk of 5%) for other indices.

## Results

### Population partitioning

Bar plots of the three independent STRUCTURE analyses carried out to evaluate population subdivision with the admixture model and without prior assumptions about time (year and epidemic period), isolate origin (plot with or without debris), and spore type giving rise to the isolate (ascospore or pycnidiospore), are provided in Fig. 2. The three cropping seasons (2009-2010, 2010-2011, 2011-2012) and the adjacent epidemic periods (end of the 2008-2009 cropping season, P01; start of the 2012-2013 cropping season, P29) are presented together. In the first set of analyses, pycnidiospore-derived subpopulations from the D+ plot were analyzed for each cropping system separately (Table S1) and then for all cropping seasons together (Table S1; Fig. 2A). The rate of change in the log probability of data between successive *K* values (Δ*K*) had a mode at *K* = 3 for the 2009-2010 season, at *K* = 2 for the 2010-2011 and 2011-2012 cropping seasons, and at *K*=7 for the overall dataset. Regardless of the epidemic period considered in each cropping season (year), it was difficult to assign individuals to any one of these *K* clusters (none of the individuals were assigned to one cluster with a posterior probability > 0.5). Instead, the posterior probability was split equally between the seven clusters (0.1426 < overall mean *P* < 0.1431), suggesting that neither the epidemic period nor the cropping season had any major effect on the structure of pycnidiospore-derived subpopulations. In the second set of analyses, pycnidiospore-derived subpopulations from the D+ and D_0_ plots were analyzed for each cropping season separately and then for all cropping seasons together (Table S1; Fig. 2B). Modes were observed at *K* = 3 for the 2009-2010 season and at *K* = 2 for the 2010-2011, and 2011-2012 cropping seasons and for the overall dataset. The posterior probability of individuals was equally distributed between the two clusters (0.498 < overall mean *P* < 0.502). Only 0.15 *%* of the individuals were assigned to one cluster with a posterior probability greater than 0.6, whatever the epidemic period, again suggesting that isolate origin (D+ or D_0_ plots) had little effect on the structure of the dataset. Finally, after the addition of ascospore-derived subpopulations to the previous dataset in the third *Structure* analysis (Table S1; Fig. 2C), a mode at *K* = 3 and evenly distributed posterior probabilities (0.332 < overall mean *P* < 0.334) were again observed. We can therefore conclude that spore type, giving rise to the isolate, had no measurable effect on population structure.

**Figure 2:**
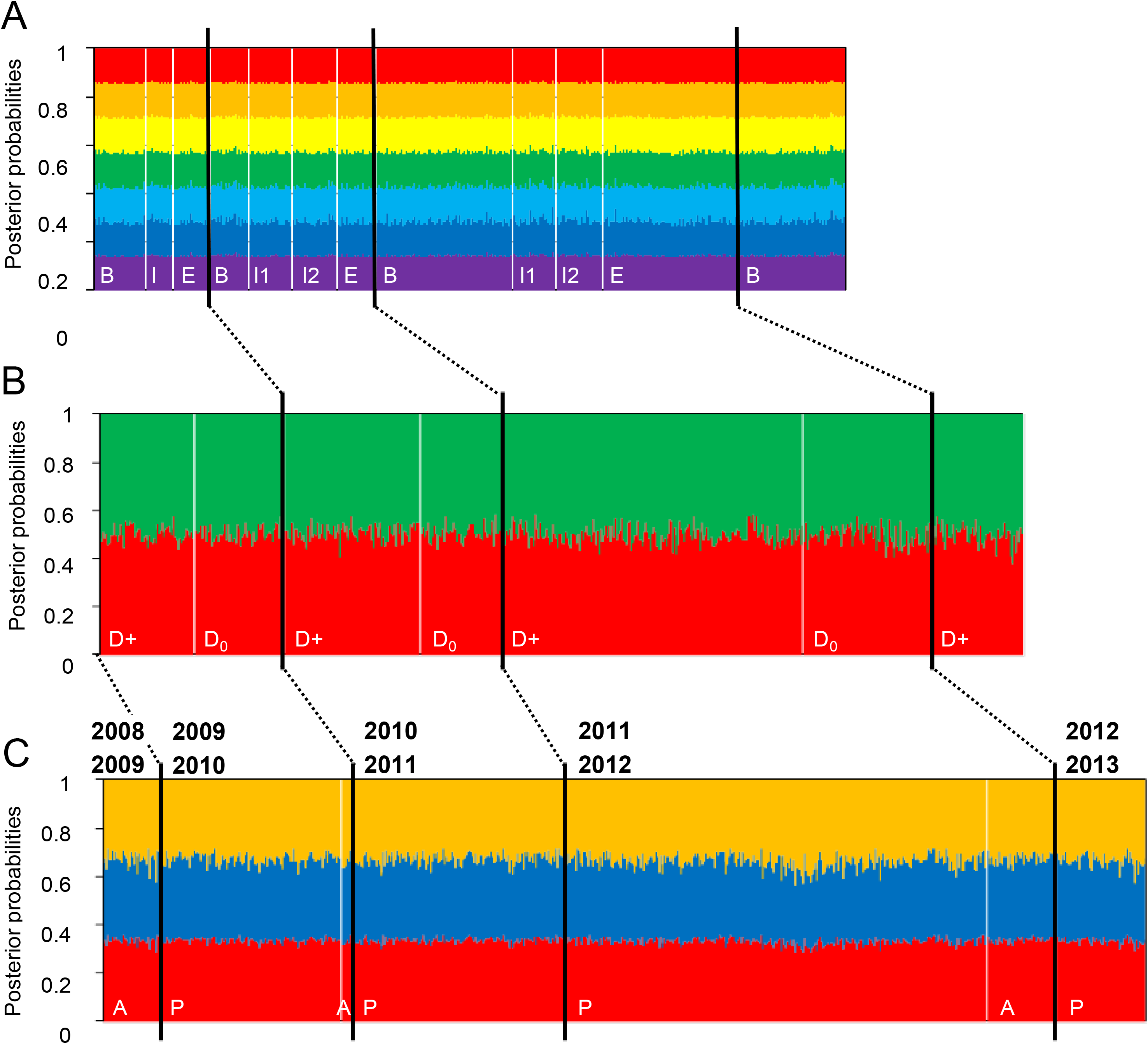
Genetic subdivision of the 794 *Z. tritici* isolates collected over the course of three successive annual epidemics (2009-2010, 2010-2011 and 2011-2012) in Thiverval-Grignon, France. Colors indicate the membership of *K* clusters for the individuals, these clusters being as defined after Structure analysis, to test three structure hypothesis. A - Effect of epidemic period (*K* = 7; 460 isolates; letters indicate the different epidemic periods (B: beginning; I1 and I2: intermediate; E: end). B - Effect of isolate origin (*K* = 2; 685 isolates; D+: plot with debris, D_0_: plot without debris). C - Spore type giving rise to the isolate (*K* = 3; 794 isolates; A: ascospores; P: pycnidiospores). For each barplot, the vertical black lines delimit the different cropping seasons and the vertical white lines delimit the second factor studied (epidemic period in A; isolate origin in B; spore type in C).

As no population structure was detected without prior assumptions, we explored the dataset with the *a priori* subpopulations defined in Fig. 1. We investigated the differentiation within and between subpopulations derived from pycnidiospores and ascospores, by calculating the pairwise *F_ST_*. The overall mean *F_ST_* was 0.016, with 22.5% of individual *F_ST_* values significant before Bonferroni correction, and 1.2% significant after this correction. Significant *F_ST_* values were obtained for populations P16, P17, P24 and P25 (Table S2).

We then examined the partitioning of genetic variation between the various factors potentially shaping the structure of *Z. tritici* populations (year and epidemic period, isolate origin, spore type), through hierarchical analyses of molecular variance (AMOVA) (Tables 1 and 2; Table S3).

A first two-factor AMOVA, assessing the effect of year (cropping season) and epidemic period nested within cropping season in each plot separately indicated that variation within epidemic periods accounted for most of the molecular variance (98-99%; *P* < 0.05; 0.012 < *F_ST_* <0.020). Variation between epidemic periods within cropping seasons accounted for only 1.7-2.1% of the molecular variance, but was nevertheless significant (*P* ≤ 0.05). Thus, the genetic variation between subpopulations was little affected by the epidemic period. We then performed a set of one-factor AMOVAs, to investigate the effect of isolate origin nested within epidemic period (Table 2). We found that, for all epidemic periods, within-plot variation accounted for most of the molecular variance (97.6-99.6%). Nevertheless, there was significant variation between plots for intermediate epidemic periods 1 and 2, even though this variation explained only a very small proportion of the molecular variance (1.9-2.4%; *P* < 0.034; 0.019 < *F_ST_* <0.024). The genetic variation of the populations was therefore slightly, but significantly, affected by isolate origin (plot D+ or D_0_), exclusively between January and April, when the number of airborne ascospores is generally lower than at the beginning and end of epidemics. A third set of AMOVA analyses (Table S3) was performed to assess the effect of spore type giving rise to the isolate on the genetic structure of subpopulations at relevant epidemic periods (end/beginning) and to test the hypothesis of temporal continuity. We compared genetic variation between the pycnidiospore-derived subpopulations collected at the end of each epidemic (P04, P21) and the ascospore-derived subpopulations present on the debris of the corresponding crop collected at the start of the next epidemic (P08, P28, respectively) in a D+ plot, and then in a D_0_ plot (P07, P27, respectively; Table S3A). Within-plot variation accounted for most of the molecular variance (96.7-100%). Between-plot variation was not significant, except for the P27-P28 comparison (3.4%; *P* < 0.05; *F_ST_* = 0.033); the difference in genetic structure between P27 and P28 was therefore small, but significant. We tested the hypothesis that early infections in D+ plots were caused principally by local inoculum resulting from a build-up of ascospores ejected from the debris of the previous wheat crop. Each ascospore-derived subpopulation collected from wheat debris at the beginning of an epidemic (P01, P08, P28) was compared with the pycnidiospore-derived subpopulations collected during the same period from D+ plots (P02, P09, P29, respectively), and from D_0_ plots (P05, P13; Table S3B). In all cases, within-population variation accounted for most of the molecular variance (89.2-100%). The differences between pycnidiospore- and ascospore-derived subpopulations were small and non-significant for both D+ plots (−0.4-3.4%; 0.116 < *P* < 0.659; −0.004 < *F_ST_*< 0.033) and for D_0_ plots (0.57% and 10.1%; 0.118 < *P* < 0.279; 0.006 < *F_ST_* < 0.101). This lack of difference in the genetic structure of the populations makes it impossible to determine, for D+ plots, whether the primary inoculum was of local or distant origin (*i.e*. originating from resident or immigrant populations).

**Table 1:**
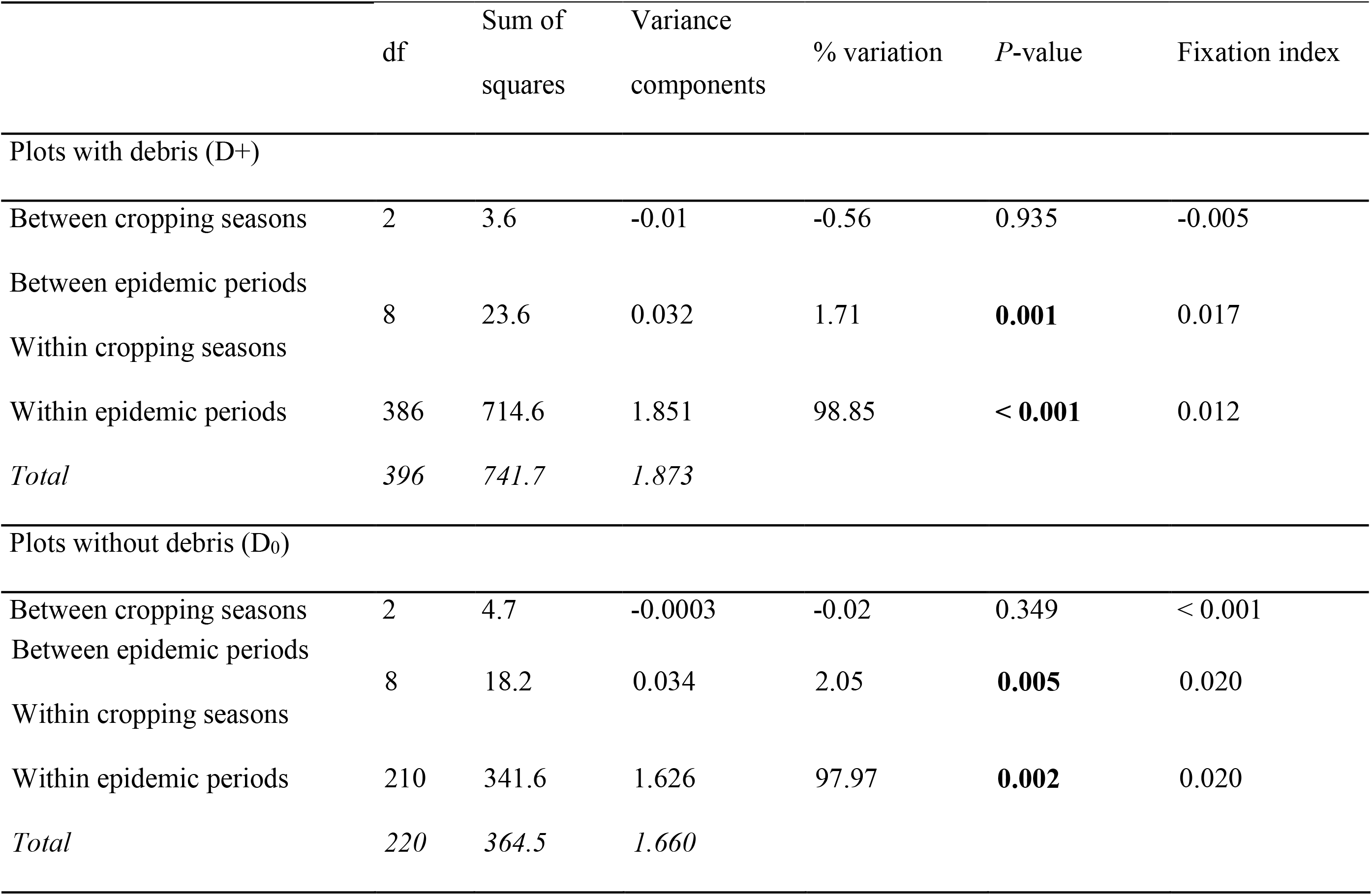
Hierarchical analysis of molecular variance (Amova) with cropping season and epidemic period nested within cropping season as grouping factors, for the D+ (upper panel) and D_0_ (lower panel) plots.

**Table 2:**
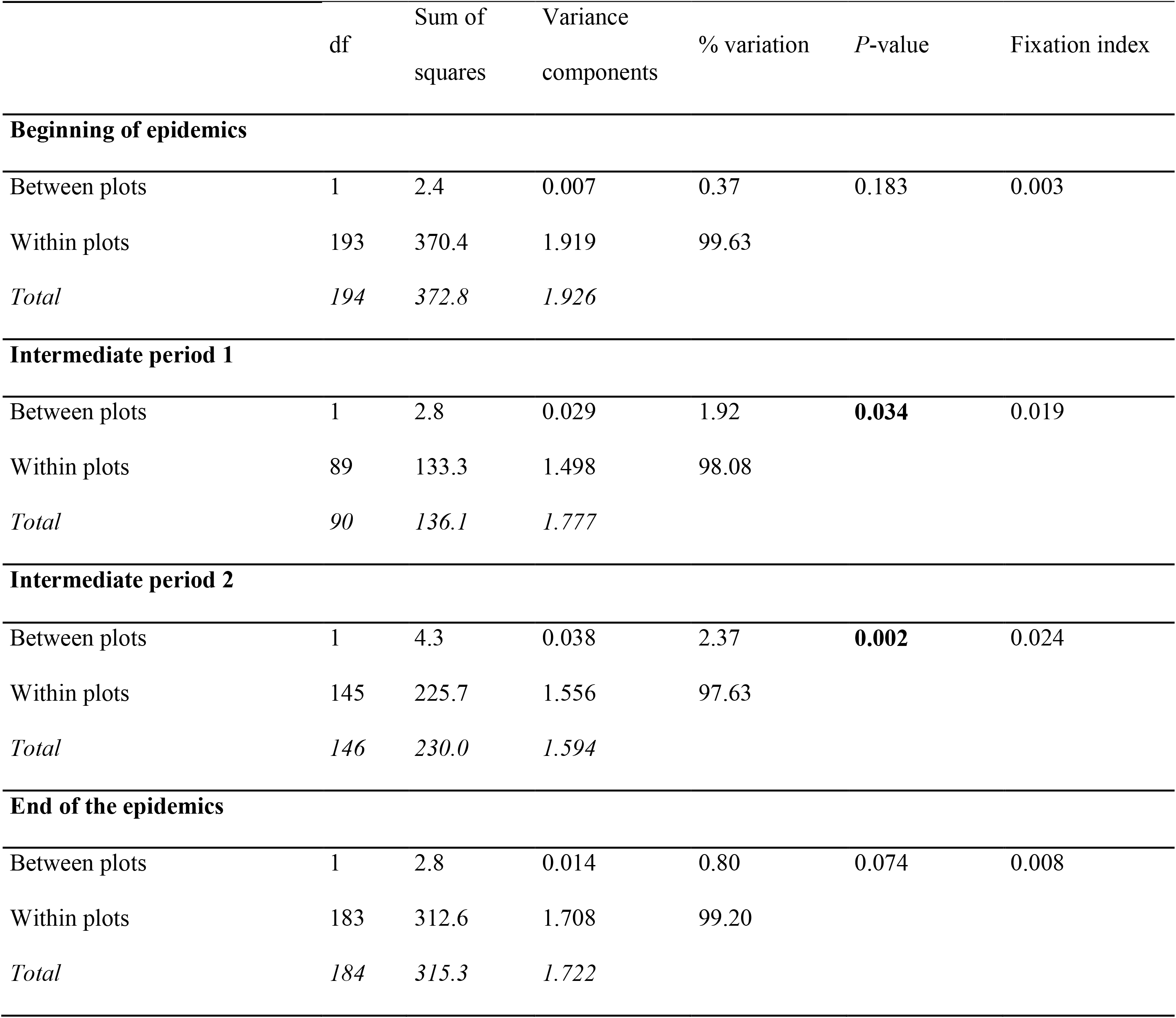
Hierarchical analysis of molecular variance (Amova), with the collection plot (D+ or D_0_) and isolate origin as grouping factors, for each epidemic period.

### Changes in population diversity over the three successive cropping seasons

The genetic variability within each of the 26 *Z. tritici* populations (Fig. 1) was the first estimated by calculating genic non-biased heterozygosity diversity, *H_E_* (Fig. 3A). *H_E_* values were of the same order of magnitude for all populations and varied between 0.38 and 0.58, with a mean at 0.46, 0.44 and 0.42 for the pycnidiospore-derived populations from D+ and D_0_ plots and for ascospore-derived subpopulations, respectively. Finally, we found no significant difference in the evolution of populations from the D+ and D_0_ plots, in a t-test for paired populations (*P* = 0.471).

**Figure 3:**
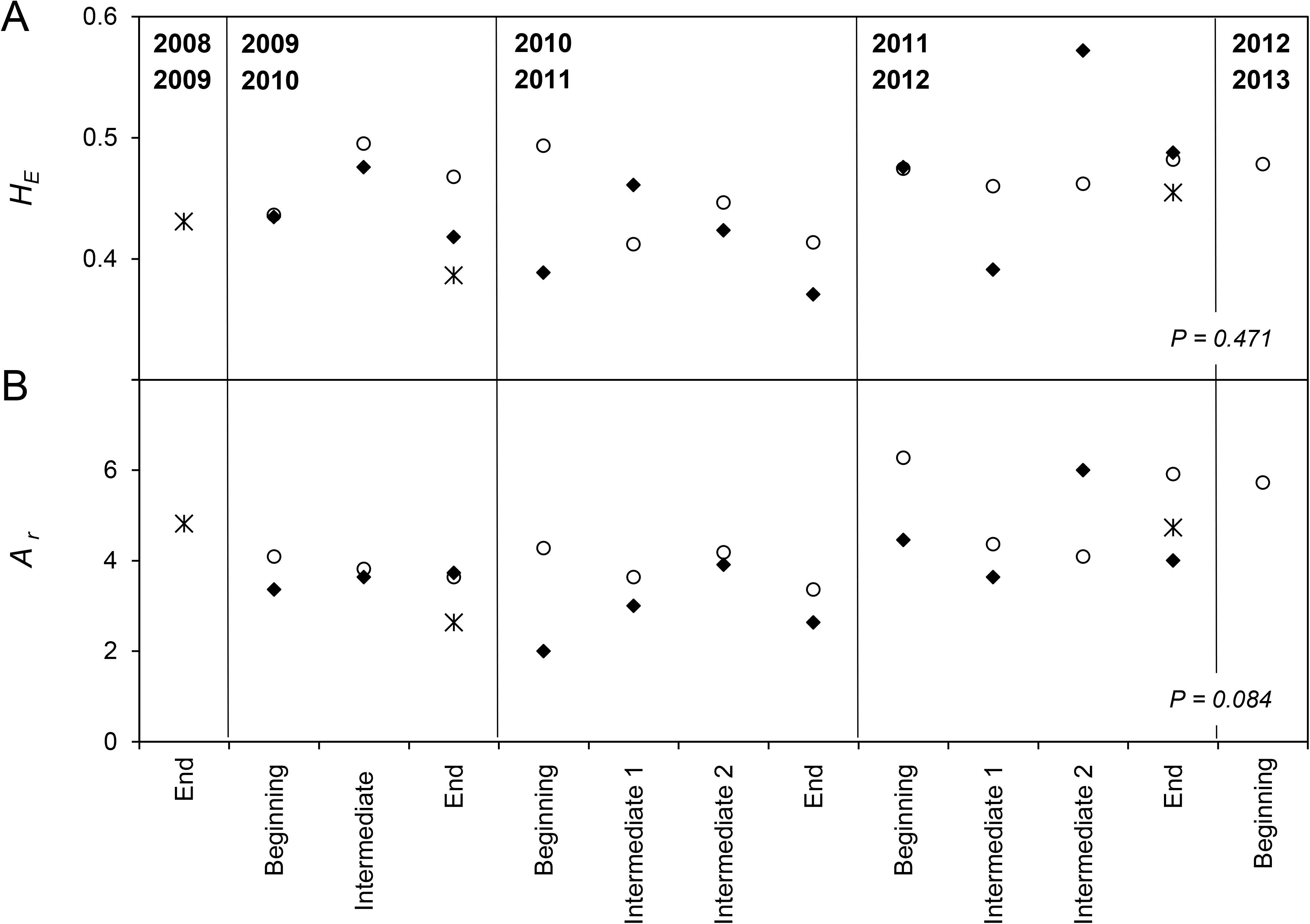
Temporal changes in the diversity of 26 *Z. tritici* populations (794 isolates; see Fig. 1) collected from wheat plots with (D+) and without (D_0_) debris, over the course of three successive annual epidemics (2009-2010, 2010-2011 and 2011-2012). A - Observed heterozygosity (*H_E_*). B - Allele richness (*A_r_*, mean number of alleles per locus). Symbols indicate the origin of the populations: white circles: pycnidiospore-derived subpopulations from D+ plots; black diamonds: pycnidiospore-derived subpopulations from D_0_ plots; crosses: ascospore-derived subpopulations.

The mean number of alleles per locus (allele richness Ar) ranged between 2 and 6.27 (mean 4.45, 3.67 and 4.06 for the D+, D_0_, and ascospore-derived subpopulations, respectively) for the three cropping seasons considered (Fig. 3B). The highest *A_r_* values were found for the 2011-2012 cropping season, which was also the season in which the number of strains collected was largest. Allele richness in the D+ plots was at least as high as that in D_0_ plots, except for the second intermediate period of the 2011-2012 cropping season, a finding not consistent with a difference in population size. No significant difference was observed between the D+ and D_0_ populations over time (*P* = 0.084).

### Occurrence of sexual reproduction in the field population of *Z. tritici*

The two mating types, MAT1-1 and MAT1-2, were found in all subpopulations (Fig. 4A) except P13 (a D_0_ pycnidiospore-derived subpopulation consisting of only five individuals of the MAT1-2 type). The MAT1-1/MAT1-2 ratio calculated for all subpopulations, including only isolates with a complete mating type (703 of the 794 isolates), was 0.95. In *χ*^2^ tests, the frequency of the two mating types was not significantly different from the expected 1:1 ratio (*P* = 0.126).

**Figure 4:**
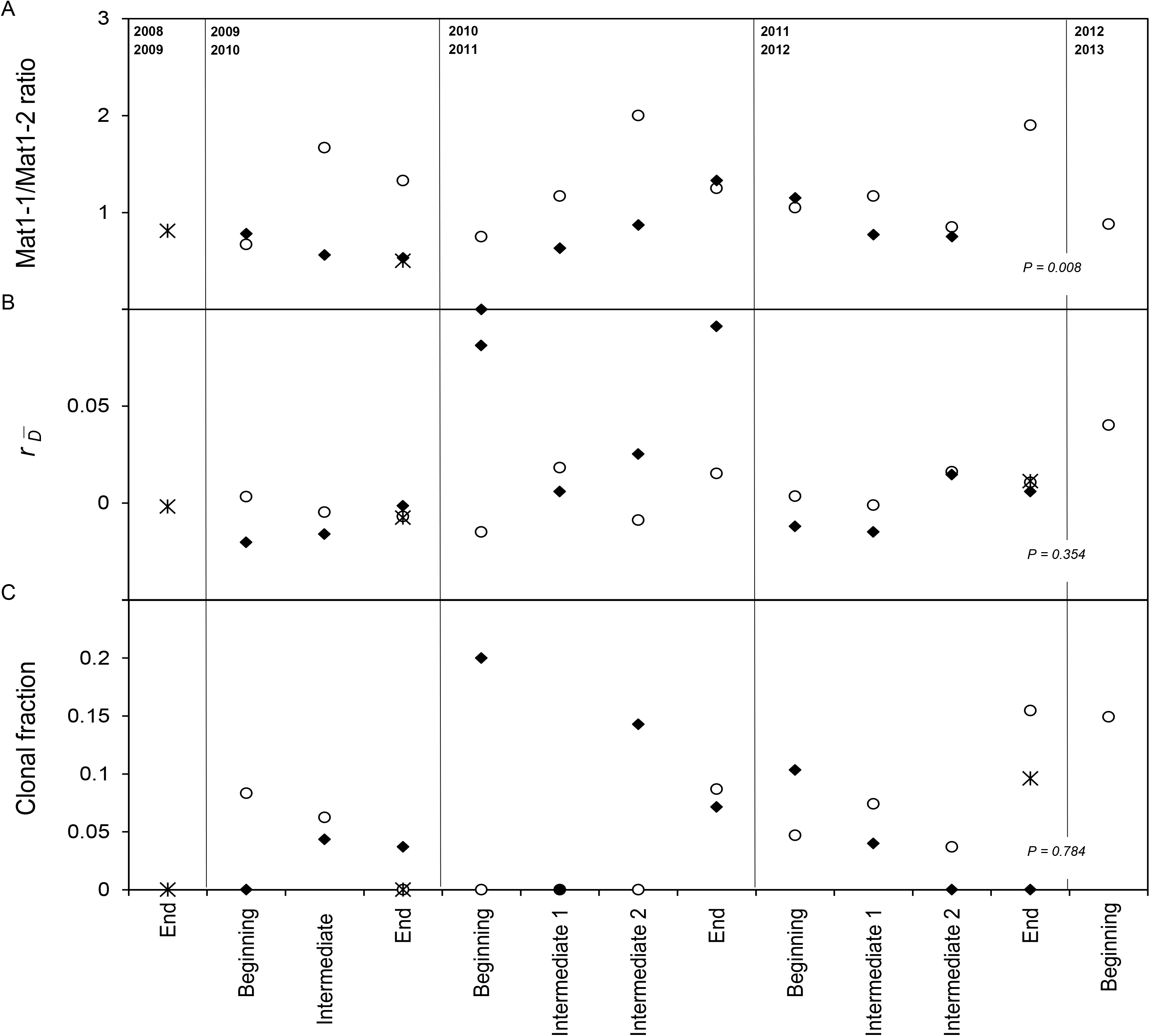
Evidence of sexual reproduction in 26 *Z. tritici* subpopulations (794 isolates; see Fig. 1) collected from wheat plots with (D+) and without (D_0_) debris, over the course of three successive annual epidemics (2009-2010, 2010-2011 and 2011-2012). A - Changes in mating-type ratio (MAT1-1/MAT1-2) over time in pycnidiospore- and ascospore-derived subpopulations. B - Changes over time in linkage disequilibrium, estimated by calculating the r_D−_ index. C - Changes over time in the clonal fraction (1-G/N) on the same plots. Symbols indicate the origin of the populations: white circles: pycnidiospore-derived subpopulations from D+ plots; black diamonds: pycnidiospore-derived subpopulations from D_0_ plots; crosses: ascospore-derived subpopulation.

We then assessed the difference in mating-type ratio between plots. The mean MAT1-1/MAT1-2 ratio ratio was 1.22 for the D+ pycnidiospore-derived subpopulations and 0.71 for the D_0_ pycnidiospore-derived subpopulations (Fig. 4A). In *χ*^2^ tests, the ratio of mating type within each plot did not differ significantly from 1:1 (P=0.407 for D+; *P* = 0.746 for D_0_). However, a significant difference in this ratio was revealed in the two plots over time (*P* = 0.008).

Finally, we assessed differences in mating-type ratio between epidemic periods. The MAT1-1/MAT1-2 ratio was lower than 1:1 in ascospore-derived subpopulations (mean ratio = 0.61; n = 92) and higher than 1:1 in pycnidiospore-derived subpopulations collected at the “end” of epidemic periods (mean ratio = 1.26; n = 605). *χ*^2^ tests revealed significant differences in mating-types distribution only between pycnidiospore- and ascospore-derived subpopulations collected at the “end” of epidemic periods (*P* = 0.006), highlighting that this ratio tends to increase over the course of the annual epidemics. This ratio was similar and close to 1:1 in “beginning” and “intermediate” pycnidiospore-derived subpopulations.

Linkage disequilibrium was always low (close to full panmixia), with 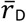 ranging from negative values to a maximum of 0.09 (P16), with a mean value of 0.006, 0.015, and 0.001 for the D+, D_0_, and ascospore-derived subpopulations, respectively (Fig. 4B). Again, no significant differences were found between the D+ and D_0_ plots (*P* = 0.354), despite 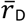 being greater at the beginning and end of the epidemic periods of the 2010-2011 cropping season in D_0_ plots.

The clonal fraction was very low over the three cropping seasons (Fig. 4C): with mean values of 0.06 for pycnidiospore-derived subpopulations from both D+ and D_0_ plots, and 0.03 for ascospore-derived subpopulations, with values ranging from a minimum of 0 (no clone found in the subpopulation) to a maximum of 0.2 (only two clones in the five-isolate subpopulation, P13). No significant difference in clonal fraction was found between the two plots (*P* = 0.784), meaning that genotypic diversity was similar. No specific pattern in clonal fraction evolution was noticed over the course of the epidemics.

## Discussion

In this study, we aimed to identify neutral structure of *Z. tritici* over the course of three successive annual epidemics at the field scale. Indeed, various inoculum sources, local or distant, may contribute to the onset and/or the development of the epidemics, with different and year-dependent timings. We hypothesized that either 1) the yearly variation in population size over epidemic courses and the regular introduction of distant “original” or “new” genotypes may induce variations in population diversity at the scale of the epidemiological cycle, or, alternatively, 2) that the demographic fluctuations endured by local populations had little effect on their diversity, with respect to the high diversity described at larger spatial scales.

We initially used an assumption-free approach for individual assignment and the testing of three putative structuring factors: time period, plot of origin and spore type giving rise to the isolate. We found no relevant neutral population structure due to any of these factors. In particular, there were no differences in the genetic structure of subpopulations collected early and late in the same season, or between the monthly samples collected from the same plot over a three-year period. These findings are consistent with the low level of between-population differentiation. These observations are also consistent with previous results showing that the genetic structure of *Z. tritici* populations was stable over time, for annual, three-year (Chen *et al*., 1994) and six-year collections (Zhan *et al*., 2001), and in space, over spatial scales ranging from meters to thousands of kilometers (Linde *et al*., 2002; Schnieder *et al*., 2001; Siah *et al*., 2018), even if variations in nested temporal smaller scales had not been checked. Accordingly, the 794 isolates collected locally constituted a single, highly diverse entity that remained stable over all epidemic stages, regardless of the demographic features of the population. This panmictic population, large and unstructured, did not appear to be affected by the putative origin of the inoculum colonizing the plots. The lack of genetic structure, resulting in similar frequencies of neutral alleles in the different subpopulations, did not allow to confirm the partially distinct origin of inoculum in the two plots which was inferred from differential adaptation to wheat cultivars established by phenotyping (Morais *et al*., 2016b). Results obtained in the current study are however not inconsistent with the previous because SSRs are supposed not “filtered” by the wheat cultivar and are so no relevant to highlight clean-cut patterns of host or local adaptation, as demonstrated by Welch *et al* (2018).

Nevertheless, the analysis of molecular variance (Amova) of populations defined *a priori* on the basis of our sampling scheme revealed that sampling time, particularly during the epidemic period, could have a slight (< 2% of molecular variance) but significant effect on the differentiation of pycnidiospore-derived subpopulations in the two plots studied. Similarly, plot of origin explained a small (< 2.5%) even if significant proportion of the molecular variance during the growing season (intermediate periods 1 and 2, i.e. between February and May, when *Z. tritici* subpopulations are thought to expand through asexual reproduction). We found that molecular variance was similar between the pycnidiospore- and ascospore-derived subpopulations collected at the end of the cropping season, with one exception (Table S3): there was weak, but significant differentiation between the pycnidiospore-derived subpopulation from the plot without debris at the end of epidemic period (P27; season 2011-2012) and the ascospore-derived subpopulation from the plot with debris collected at the beginning of the next season (P28; 2012-2013). This effect was not detected in previous studies of single strains collected yearly (Chen *et al*., 1994; Zhan *et al*., 2001). The differences could be an artefact, due for instance to the sensitivity level of the AMOVA test used or to the sample size.

Genetic diversity was high to very high in all subpopulations, and stable over the three cropping seasons (variation over time non-significant). The orders of magnitude of expected heterozygosity (H_E_), allele richness (A_r_) and the proportion of multilocus haplotypes were consistent, although somewhat smaller, with the values obtained with SSRs and RFLPs over larger geographic scales for Israel, Swiss, USA, Tunisian, French and Canadian populations (Linde *et al*., 2002; El Chartouni *et al*., 2011; Boukef *et al*., 2012; Razavi & Hughes, 2004).

Pycnidiospore- and ascospore-derived subpopulations did not differ in diversity, whereas lower levels of clonal fraction might have been expected after sexual recombination, which is thought to occur in winter. No significant difference in the evolution of genetic and genotypic diversity was found between the two plots, so the larger amount of local primary inoculum produced in the plots containing debris had little or no effect on population diversity. This contrasts with the selection thought to be exerted on some loci during the epidemic period by cultivar (Morais *et al*., 2016b), temperature (Suffert *et al*., 2015), or fungicides (Hayes *et al*., 2016) or with the putative counter-selection during the interepidemic period (Suffert *et al*., 2018). Our findings are not inconsistent with such selective dynamics as we focused actually on neutral diversity. Population differentiation at quantitative, adaptive traits is not directly linked with neutral genetic diversity or differentiation; even if neutral genetic markers have great potential for investigating processes such as gene flow, migration or dispersal, this type of genetic variation tells little about the adaptive or evolutionary potential of a population (Holderegger *et al*., 2006), except in situations where clear founder effects are demonstrated (Jürgens *et al*, 2005). Other markers could be developed from recently described genes determining *Z. tritici* adaptation, for instance adaptation to the Stb6 resistance gene present in many French cultivars (Zhong *et al*., 2017; Brunner & MacDonald, 2018) or to DMI fungicides largely used in Western Europe (Cools *et al*., 2013), and would help to clarify this issue. Such selected markers would be particularly relevant when the frequency of these genes is still moderate (e.g. range from 5 % to 25 %) and not homogeneously distributed at a small spatial scale; this may be the case for a virulence corresponding to a resistance very recently overcome or the case of a fungicide resistance recently appeared.

The presence of both idiomorphs, MAT1-1 and MAT1-2, in all populations, albeit in variable proportions, confirmed that sexual reproduction could occur all year round (Duvivier, 2015). Linkage disequilibrium remained very low over time, at values close to full panmixia, regardless of primary inoculum origin. Similarly, the clonal fraction was always very low, regardless of sampling time and inoculum origin. For example, the 2009-2010 cropping season was characterized by more intense ascospore production in the plot with debris than during the other two cropping seasons studied (Morais *et al*., 2016a), but this had little effect on the patterns of diversity indices over the cropping season. Overall, these results are consistent with recombination occurring regularly, due to sexual reproduction over a long period of the year, but impossible to pin down with these data, which are, again, of a similar order of magnitude to data already reported for this fungal species at a larger geographic scale.

The overall ratio of the two mating types in the whole population did not differ significantly from the expected 1:1 ratio, consistent with previous results (Zhan *et al*., 2002; Siah *et al*., 2010). However, an interesting finding is that ratios of mating-types change over time. Frequency of MAT1-1 strains increased over the course of the epidemics, suggesting that MAT1-2 exhibited a competitive disadvantage and was counter-selected during the cultural season through the asexual cycles. This hypothesis is consistent with the average difference in pathogenicity between the two mating-types (MAT1-1 isolates having 14-22% greater pathogenicity than MAT1-2 isolates) previously reported by Zhan *et al* (2007), and with selective dynamics over the course of a growing season highlighted by Suffert *et al*. (2016; 2018). This is the first evidence of this property expressed and having epidemiological consequences under field conditions. This finding of a significant change in mating type frequencies at short term is an exciting observation that deserves further exploration *via in planta* studies.

Finally, local *Z. tritici* populations displayed high levels of genetic uniformity between years, epidemic periods and plots, which is consistent with the results obtained by Chen *et al*. (1994), Zhan *et al*. (2002) and El Chartouni *et al*. (2012) but was never established at a fine temporal scale. Our demogenetic analysis of a local population thought to result from different evolution trajectories through three successive sequences of epidemic and interepidemic periods highlights only sporadic and small effects of the type of inoculum, depending on epidemic period and year. Analogous hypotheses about the respective contributions of sexual/asexual primary/secondary inoculum were tested by Cortesi & Milgroom (2001) and Shah *et al*. (2001). Such studies, which remain uncommon at fine spatiotemporal scale, show how population genetics can be used to address epidemiological questions essential for sound disease management (Milgroom & Peever, 2003) but also what are its limits and opportunities for further improvement.

## Acknowledgments

The authors thank Angélique Gautier (INRA BIOGER) and Christophe Montagnier (INRA Experimental Unit, Thiverval-Grignon) for technical assistance. This study was supported by a grant from the European Union Seventh Framework Program (Grant Agreement no. 261752, PLANTFOODSEC project) and a grant from the European Union Horizon Framework 2020 Program (Grant Agreement no. 634179, EMPHASIS project).

